# *Cytocipher* determines significantly different populations of cells in single cell RNA-seq data

**DOI:** 10.1101/2022.08.12.503759

**Authors:** Brad Balderson, Michael Piper, Stefan Thor, Mikael Boden

## Abstract

Identification of cell types using single cell RNA-seq (scRNA-seq) is revolutionising the study of multicellular organisms. However, typical scRNA-seq analysis often involves post hoc manual curation to ensure clusters are transcriptionally distinct, which is time-consuming, error-prone, and irreproducible. To overcome these obstacles, we developed *Cytocipher*, a bioinformatics method and *scverse* compatible software package that statistically determines significant clusters. Application of *Cytocipher* to normal tissue, development, disease, and large-scale atlas data reveals the broad applicability and power of *Cytocipher* to generate biological insights in numerous contexts. This included the identification of cell types not previously described in the datasets analyzed, such as CD8+ T cell subtypes in human peripheral blood mononuclear cells; cell lineage intermediate states during mouse pancreas development; and subpopulations of luminal epithelial cells over-represented in prostate cancer. *Cytocipher* also scales to large datasets with high test performance, as shown by application to the Tabula Sapiens Atlas representing >480,000 cells.

*Cytocipher* is a novel and generalisable method that statistically determines transcriptionally distinct and programmatically reproducible clusters from single cell data. *Cytocipher* is available at https://github.com/BradBalderson/Cytocipher.

## Introduction

ScRNA-seq has transformed the study of complex tissues, allowing data-driven characterisation of cell types through the bioinformatics analysis of genome-wide gene expression measurements across thousands to millions of cells^1^. The current standard bioinformatics analysis pipeline consists of normalisation of read counts, gene/feature selection, dimensionality reduction, nearest-neighbour graph construction, and finally clustering and visualisation based upon the nearest-neighbour graph^1^.

The most widely used approach for clustering is the Leiden algorithm^2^, which considers the nearest-neighbour graph as the primary representation of the single cell data^2^, and thereby does not consider the original gene expression measurements. To determine if clusters of cells represent transcriptionally distinct populations post hoc differential expression analysis is performed. Based on visual assessment of the resulting differentially expressed genes, Leiden clustering is adjusted or clusters manually altered if automated clustering cannot align with observed transcriptionally distinct cells.

The manual merging process, in addition to being irreproducible and non-quantifiable, can be impractical. For example, 1) when cell populations do not have any one single marker gene that differentiates one cluster from others but may be unique due to a *combination* of genes, which is difficult to visualise, or 2) when the presence of tens to hundreds of cell types make the manual merging process time consuming and prone to human error.

Ensuring that scRNA-seq clusters represent transcriptionally distinct populations of cells is a currently unaddressed issue in the unsupervised analysis of scRNA-seq data. To address this fundamental problem, we present *Cytocipher*, a *scverse* compatible package that integrates within the Python *Scanpy*^*1*^ ecosystem for single cell analysis. *Cytocipher* implements two novel algorithms that address the aforementioned problems; 1) *code-scoring* - a novel per-cell enrichment scoring method that is more sensitive at scoring unique *combinations* of genes within individual cells when compared with current methods, and 2) *cluster-merge* - a statistical algorithm that begins with over-clustered single cell data and iteratively tests clusters for significant transcriptional difference, merging clusters that are not mutually significantly different. The resulting output is significantly transcriptionally different single cell populations, the marker genes for each population (Box 1), and per-cell enrichment scores for marker gene sets (Box 1).

### Box 1.

Glossary

- **Clustering:** the algorithmic process for grouping observations (e.g. cells) based on a given metric of similarity.
- **Over-clustering:** clustering that results in a set of clusters where *two* or more of the clusters do not contain observations with features that are distinctly different (e.g. cells in separate clusters that express the same genes).
- **Under-clustering:** clustering that results in a set of clusters where *one* or more of the clusters contain observations that do not share a distinctive feature or combination of features (e.g. cells grouped that do not express any common genes).
- **Enrichment scoring:** Quantitative scoring of the collective expression of a gene set in a gene expression profile.
- **Marker genes:** given a set of clusters for single cells, the set of genes for each cluster which maximally demarcate a given cluster from all others.
- **Cross-scoring:** Given a set of marker genes for a set of clusters, the case where performing enrichment scoring for all cells and marker gene sets results in cells belonging to a given cluster receiving high enrichment scores for marker genes of a different cluster.

To probe the versatility of *Cytocipher* we applied it to synthetic data and a range of biological samples - healthy adult tissue, development tissue, diseased tissue, and multitissue atlas scale data. Each application resulted in the identification of biological insights not previously described, underscoring the power of *Cytocipher* as a universal method for improving single cell population detection on previously analysed or newly generated scRNA-seq data. *Cytocipher* also scales to large datasets with high test performance, as shown by application to the Tabula Sapiens Atlas^3^ representing >480,000 cells.

The ongoing Atlas scale single cell data generation^3^ is producing increasingly complex and heterogeneous single cell datasets, underscoring the importance of a statistically rigorous test to ensure cell cluster determination represents transcriptional distinction. *Cytocipher* represents the first analysis method that frames transcriptionally distinct single cell population detection as a quantifiable statistical test, thereby representing an important advance in the analysis of scRNA-seq data.

## Methods

### Cytocipher code-scoring

*Cytocipher code-scoring* is a per-cell enrichment scoring method (Box 1) that ensures that the unique *combination* of marker genes (Box 1) are expressed in cells which receive positive scores. This contrasts with existing methods, which utilise an additive score that can positively score cells even if only a single marker gene is expressed^1,4,5^. Log normalised single cell gene expression and tentative cluster labels are required as input (Figure 1A1). If input of marker genes is not provided these are automatically determined (Box 1, Figure 1A2). By default, the marker genes per cluster are gene sets of up to 5 differentially upregulated genes (Benjamini-Hochberg adjusted p-value < .05) determined by ranking genes using a one-versus-rest mode of comparison with Welch’s t-statistic. With marker genes per cluster (*G*, Figure 1A2), negative gene sets per cluster are determined (Figure 1A3). Given the set of clusters (*C*), we define the negative gene sets (*N*_*i*_) for a given cluster *i* with marker gene set *G*_*i*_ as the set of gene set differences between *G*_*i*_ and each other marker gene set *G*_*j*_ for which the intersection of *G*_*i*_ and *G*_*j*_ is greater than or equal to a minimum intersection count (Equation 1).

**Figure 1.**
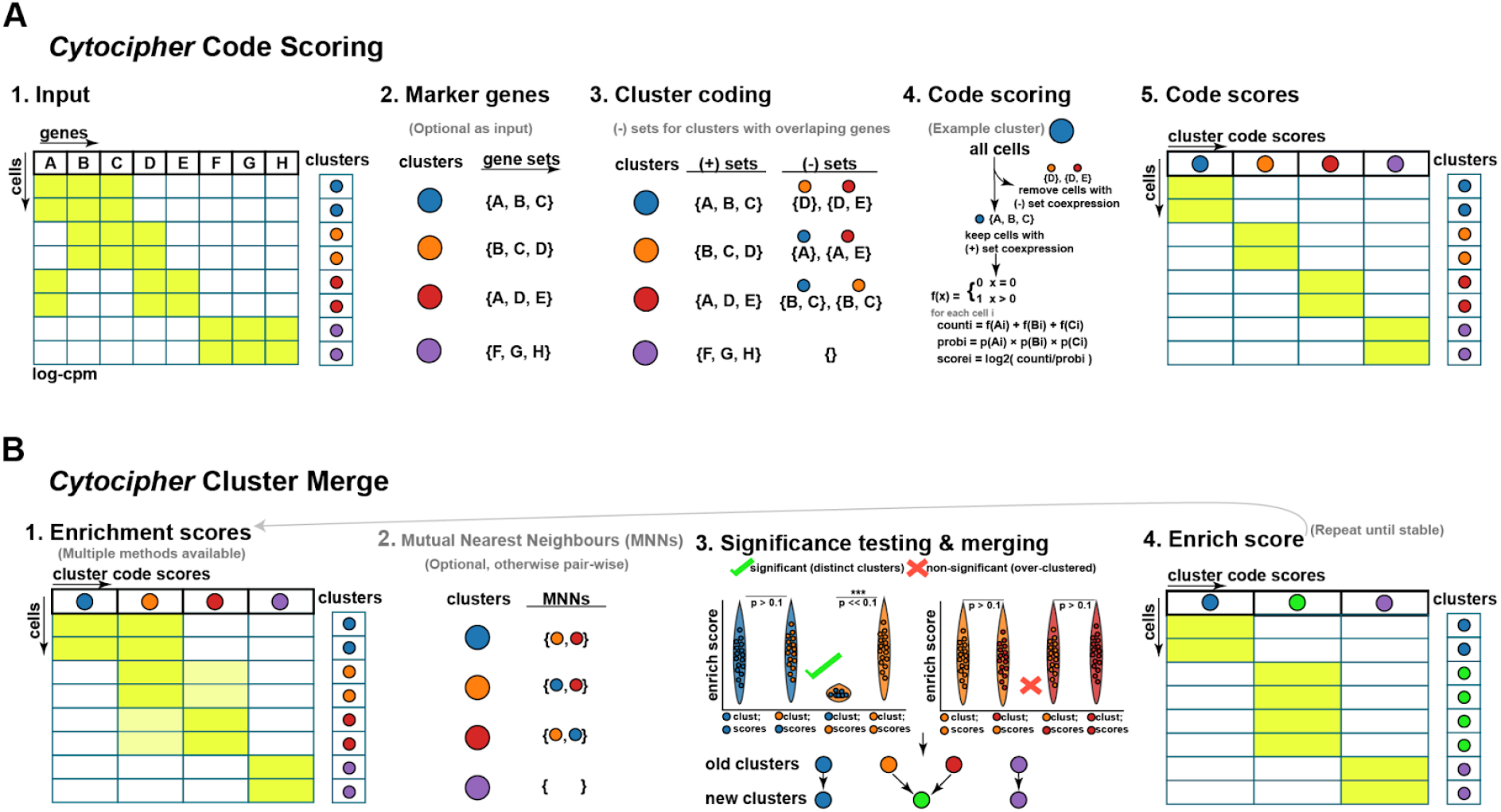
Overview of *Cytocipher* for cluster specific *code-scoring* and significant cluster analysis with *cluster-merge*. **A**. Schematic illustration of the *Cytocipher code-scoring* method. Briefly, 1) Gene expression and cluster annotation are provided as input; 2) determination of marker genes is performed; 3) genes which are positive indicators versus negative indicators of cluster membership are determined through comparison of cluster marker genes; 4) for a given cluster, cells co-expressing the negative gene sets are filtered, while cells co-expressing the positive gene set are kept, followed by co-expression scoring; 5) repetition of 1-4 for each cluster yields a diagnostic heatmap scoring each cell for membership of each cluster. **B**. Schematic illustration of *Cytocipher cluster-merge*, which performs a test of significance difference between cluster pairs, merging those which are *mutually* non-significantly different. Briefly, 1) per cell enrichment scores are determine using the process in A; 2) where a large number of clusters are present, mutual nearest neighbours can determine cluster pairs for comparison; 3) cluster scores are compared with a statistical test to determine significant versus non-significant clusters, with non-significant cluster pairs being merged to create new cluster labels; 4) Enrichment scores are redetermined based on the new cluster labels as an output diagnostic, and the process can be repeated with the new cluster labels until convergence.

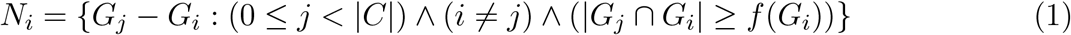

The minimum intersection size function, *f* (Equation 2), acts as a winsorization for small gene set sizes, such that if the marker gene set for cluster *i* (*G*_*i*_) has a size of 1 or 2 genes, all genes must intersect with the marker gene set of cluster *j* (*G*_*j*_) for the marker gene set difference to be added to the negative gene set (*N*_*i*_). If a marker gene set has 2 or more genes, all genes except for one must overlap with the marker gene set of another cluster *j* (*G*_*j*_) for the marker set difference to be added to the negative gene set (*N*_*i*_).

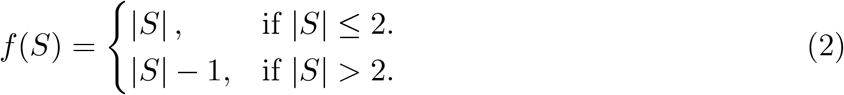

Letting Ω represent the cell by cluster code-score matrix of size *T ×* |*C*|, we define the *code-scoring* for cluster *i* and cell *κ* (Ω_*κ,i*_) as in Equation 3 (Figure 1A4-5).

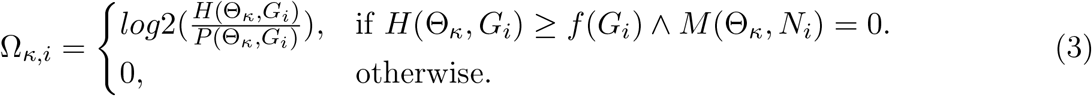

Θ represents a cell by gene expression matrix of size *T × F*, such that Θ_*κ,g*_ represents the expression of gene *g* in cell *κ. H*(Θ_*κ*_, *G*_*i*_) is the number of genes in *G*_*i*_ expressed in cell *κ* (Equation 4), *M* (Θ_*κ*_, *N*_*i*_) is the number of negative gene sets co-expressed in cell *κ* (Equation 5) and *P* (Θ_*κ*_, *G*_*i*_) is the probability of jointly expressing genes in *G*_*i*_ at the observed expression level in cell *κ* across all cells (Equation 6).

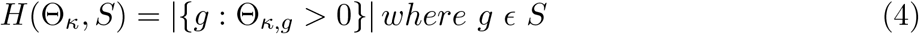

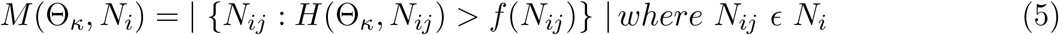

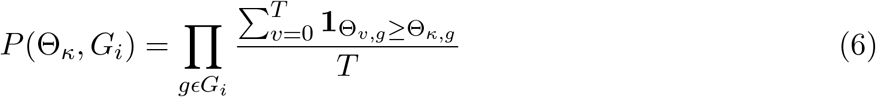

An additional enrichment scoring method, *coexpr-scoring* (Ψ_*κ,i*_, Equation 7), is defined without consideration of negative gene sets (*N*_*i*_) to assess the effect of negative gene sets on cluster scoring.

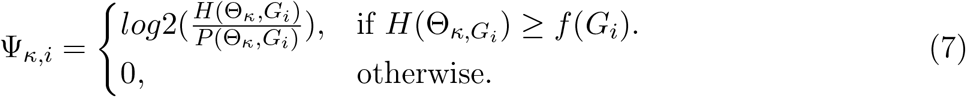

Intuitively, the greater the number of genes co-expressed and the greater the expression level, the higher the total code-score and coexpr-score; with *code-scoring* ensuring non-intersecting marker genes are not expressed in similar clusters.

### Cytocipher cluster-merge

*Cluster-merge* requires tentative cluster labels for each cell (*z*) and Ω. Ω can also be substituted for other enrichment score matrices, such as Ψ (Figure 1B1).

For a given pair of clusters *i* and *j*, two independent Student’s t-tests on enrichment scores determine if they are significantly different (Figure 1B3). Let the notation Ω_*j∼i*_ indicate enrichment scores for cluster *i* across all cells that belong to cluster *j*. Given the function *t* below represents Student’s t-value calculation, and the function *p* below indicates the probability of observing a given t-value using a two-sided test; we define a significantly different pair of clusters (*s*_*ij*_) as in Equation 10.

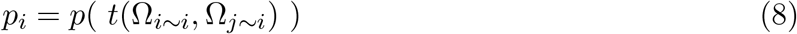

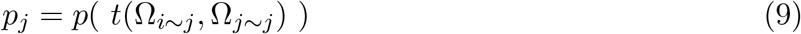

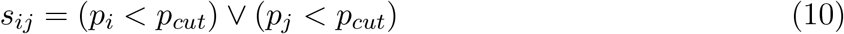

*p*_*i*_ is the p-value that cluster *i* and cluster *j* cells have the same cluster *i* enrichment scores, while *p*_*j*_ is the p-value the cluster pair cells have the same cluster *j* enrichment scores (p-value adjustment with Benjamini-Hochberg) (Figure 1B3). Clusters are significantly different (*s*_*ij*_) if either the *p*_*i*_ or *p*_*j*_ are below a significance threshold (*p*_*cut*_). If the cluster pairs are *mutually*

*non-significantly different* (*s*_*ij*_ = *False*), cells from the cluster pairs are merged to create a new set of cluster labels (*z*_*new*_) (Figure 1B3).

By default, *p*_*cut*_ is taken as 0.01 and each cluster is compared against all others. In the case of many clusters mutual nearest neighbours (MNNs) can be compared to reduce run time (Figure 1B2). Cluster MNNs are cluster pairs with overlapping *φ*-nearest neighbours using the Euclidean distance metric on a |*C*| *×* |*C*| matrix *ξ*; where *ξ*_*ij*_ is the mean cluster *j* enrichment score in the cells of cluster *i*.

Marker genes, enrichment scoring, and cluster merging can be performed iteratively until convergence (*z*=*z*_*new*_) (Figure 1B4). In the initial data tested, convergence occurred after a single iteration, prompting this as the default setting.

### Enrichment score summarisation

High false positive rates and inflated power due to treating each cell as an independent observation has been previously observed in the context of single cell differential expression analysis, and is addressed by ‘pseudobulking’^6,7^. We systematically tested different methods for summarising enrichment scores Ω_*i∼i*_ and Ω_*j∼i*_ (if scoring for cluster *i*) prior to significance testing with the goal of maintaining test performance while reducing inflated p-values and correlation of significance with the total number of cells in a cluster pair. Three different methods were compared using the 3,000 human peripheral blood mononuclear cells (PBMCs) scRNA-seq data; determine *k* evenly spaced quantiles (K-Quantiles), taking the mean of enrichment scores falling into *k* evenly spaced quantile bins (K-Bins), and clustering enrichment scores into *k* groups with K-Means clustering and averaging the enrichment scores in each group (K-Means). The PBMCs were first over-clustered (see Methods below) prior to applying each summarisation method with *k* =15. The original PBMC annotations were used as a ground-truth, such that cluster pairs falling within a cell type were considered non-significantly different and pairs from different cell types were considered significantly different. Test performance was measured by the area under the curve (AUC) of the receiver operating characteristic (ROC, AUROC). Cell abundance bias was measured using Spearman’s correlation with -log10(p-value) and log2(cluster pair cell count). P-value cut-offs to discriminate significant versus non-significant cluster pairs using *Cytocipher* were from 0 to 1 in increments of 2^−*i*^ for each integer *i* in range 0 ≤ *I* ≤ 100. Parameter sensitivity analysis was then performed varying *k* and measuring the cell abundance bias and test performance for each summarisation method. The above tests prompted usage of the K-Quantiles method with k=15 as the default method for enrichment score summarisation prior to cluster pair significance testing with *Cytocipher*.

### Simulation study

*Splatter*^*8*^ v1.20 was used to simulate initial scRNA-seq read counts followed by post hoc alteration of counts to simulate groups that express a unique *combination* of genes, rather than a single gene marker. Simulation parameters were estimated from the PBMC gene *×* cell count matrix with *splatEstimate*. 10,000 genes and 3,000 cells were then simulated from the learnt parameters. 8 groups of random cells were assigned a unique random set of 3 to 7 genes drawn from 15 genes to become differentially expressed (DE).

DE was simulated by multiplying gene counts within a cluster by factor *φ* drawn from Gaussian distribution *N* (*µ*_*φ*_ = 6, *δ*_*φ*_ = 1). 0 count cells were set to the median non-zero count value prior. 0 count cells set to the median was determined by sampling proportion *τ* from Gaussian *N* (*µ*_*τ*_ = 0.85, *δ*_*τ*_ = 0.05). Default preprocessing and normalisation with *Scanpy*^*1*^ v1.8.2 was subsequently performed on the simulated counts.

Enrichment scoring was performed for each method with default parameters. For *Scanpy* - scoring, we refer to the sc.tl.score_genes, a re-implementation of the scoring approach provided by the Seurat package^9^.

Cells with positive scores for a given group were classified as belonging to the group, and cells with 0 or less scores for a given method as outside of the group. Enrichment score-based annotations were quantitatively benchmarked against the original labels for each group using accuracy, precision, recall, and F1 score calculated with *Sklearn*^*10*^.

### E18.5 mouse hypothalamus study

Single cell count matrices, cell annotations, and UMAP coordinates were obtained from the original study^11^. Gene expression was normalised using log1p counts normalised to total cell counts with the median read counts observed across all cells in *Scanpy*. Enrichment methods were applied using default settings on the neuropeptidergic-dopaminergic neuron subtypes defined in the original study with the annotated marker genes. Quantitative benchmarking of enrichment methods was performed equivalently to the simulated data above.

### Human 3K peripheral blood mononuclear cell (PBMC) study

The human 3K PBMC single cell count matrices were downloaded and preprocessed as per the *Scanpy* PBMC 3K tutorial^1^, except for total count and cell mitochondrial percentage regression. Leiden clustering was performed at resolution 0.7 producing the same 8 clusters in the vignette. For the *Cytocipher cluster-merge* analysis, the PBMC scRNA-seq was first over-clustered using Leiden resolution 2.0 resulting in 19 clusters. *Cytocipher code-scoring* was performed with default settings. *Cytocipher cluster-merge* was also performed with default settings.

### Mouse E15.5 pancreas development study

The mouse E15.5 pancreas development scRNA-seq spliced and unspliced read count matrices and cell annotations were downloaded as per the *Scvelo*^*12*^ v0.2.4 *RNA Velocity Basics* tutorial. Genes with a minimum shared counts of intronic versus exonic reads of 4 were filtered prior to log1p and total library size normalisation. The velocity UMAP embedding was then reproduced from the original *Scvelo* publication using the same parameters and commands detailed in the *Scvelo* tutorial aforementioned. Cells were over-clustered using a Leiden resolution of 3.5. *Cytocipher cluster-merge* was performed with default parameters and cluster pairs were considered significantly different at an adjusted p-value cutoff of 0.045.

### Prostate cancer study

The *Anndata* h5ad file containing the normalised gene expression, cell annotations, and UMAP was obtained from the original study^13^. The cell-cell neighbourhood graph was constructed using 30 nearest neighbours and over-clustered with a Leiden resolution of 4.0 to create 47 tentative clusters. Marker genes per tentative cluster were determined as a maximum of 4 genes and minimum of 1 gene ranked by log-FC using a one-versus-rest mode of comparison for each cluster for which Welch’s t-statistic was above 14. *Cytocipher cluster-merge* was otherwise applied with default values, with an adjusted p-value cutoff of 0.0325 used to determine significantly different cluster pairs.

Differentially abundant subpopulations of cells between cancer and normal cells were determined using the *Milopy* implementation of the *Milo* method^14^. Parameters used were the same as *Milopy* tutorial on mouse gastrulation scRNA-seq. Exceptions were the usage of *∼group* as the linear model inputted to *milo.DA_nhoods* function in order to call differentially abundant cell populations between cancer and normal samples in the prostate cancer data.

### Tabula Sapiens study

The *Anndata* h5ad file containing all preprocessed data was obtained from the original study^3^. In all applications mentioned, default parameters were used with 15 CPUs on the highly variable genes.

To generate ground-truth labels of significant versus non-significant cluster pairs, all cell types with more than 20 cells were split into 3 random subgroups. Cell types with less than 20 cells were not split. Cluster pairs were labelled as significantly different in this ground-truth if they were sampled from different cell types and non-significantly different if sampled from the same cell type. *Cytocipher cluster-merge* was initially applied on the artificial subgroups with an adjusted p-value cutoff of 0.5 to examine the topmost over-merged clusters. ROC curves were determined using the ground-truth cluster pair labels compared against significant versus non-significant pairs determined at the respective p-value cutoff outputted by *Cytocipher cluster-merge*. P-value cutoffs used were the same as detailed in the section *Enrichment Score Summarisation*. Memory usage was measured as the change in peak random-access memory (ΔRAM). Each artificial subgroup was then downsampled to a maximum of 15 cells resulting in 7,385 cells.

*Cytocipher cluster-merge* was compared with *Sc-SHC* on a small subset of 500 cells representing 12 randomly sampled cell types from the 177 total cell types in the downsampled 7,385 cell data. Both methods were run with default parameters on the highly variable genes. *Sc-SHC* was run on the ambient RNA corrected *DecontX*^*15*^ read counts.

### Data Availability

All data analysed are publicly available. The E18.5 hypothalamus neuronal subtype data is available in the Gene Expression Omnibus (GEO) database at accession GSE154995 with the cell annotations and UMAP coordinates available in the supplemental data of the original paper^11^. The E15.5 mouse pancreas data^16^is available in GEO at accession GSE132188. The processed data was downloaded using *Scvelo* with the scv.datasets.pancreas() function. The human PBMC 3K data are hosted by 10X Genomics, and were downloaded via wget http://cf.10xgenomics.com/samples/cell-exp/1.1.0/pmbc3k/pbmc3k_filtered_gene_bc_matricies.tar.gz. The prostate cancer data was downloaded from http://www.prostatecellatlas.org/ with wget https://cellgeni.cog.sanger.ac.uk/prostatecellatlas/prostate_portal_300921.h5ad. The Tabula Sapiens data is available from figshare https://figshare.com/projects/Tabula_Sapiens/100973 and was downloaded via wget https://figshare.com/ndownloader/files/34702114.

### Code Availability

*Cytocipher* is available at https://github.com/BradBalderson/Cytocipher. The tutorial includes full code to reproduce the mouse pancreas analysis.

## Results

### *Cytocipher code-scoring* outperforms existing methods for binary classification task on simulated data

To test *Cytocipher code-scoring* for specifically scoring cells expressing a unique *combination* of genes, we simulated scRNA-seq data where clusters are defined by unique gene combinations, rather than single marker genes (Figure 2A-B; see Methods for simulation details). As an example, cluster 7 in our simulation expresses the unique gene combination 3094, 5076, 6481, 702, and 7274, but none of these individual genes are specifically expressed only in cluster 7. We subsequently scored each cell for the cluster marker genes using four different approaches: *Cytocipher code-scoring, Cytocipher coexpr-scoring* (*code-scoring* without negative set filtering), *Giotto PAGE*^*5*^, and *Scanpy-score*^*1*^ (Figure 2C). Qualitatively, in every cluster *code-scoring* specifically highlighted the cells belonging to the relevant clusters, even in the difficult cases where no specific marker gene demarcated the cluster. The clearest examples of this are clusters 4 and 7. In the former case, all other methods scored cells in cluster 0 and 4 when attempting to score for cluster 4 (see cross-scoring, Box 1). In the latter case cross-scoring was observed with cluster 2 when scoring for cluster 7.

**Figure 2.**
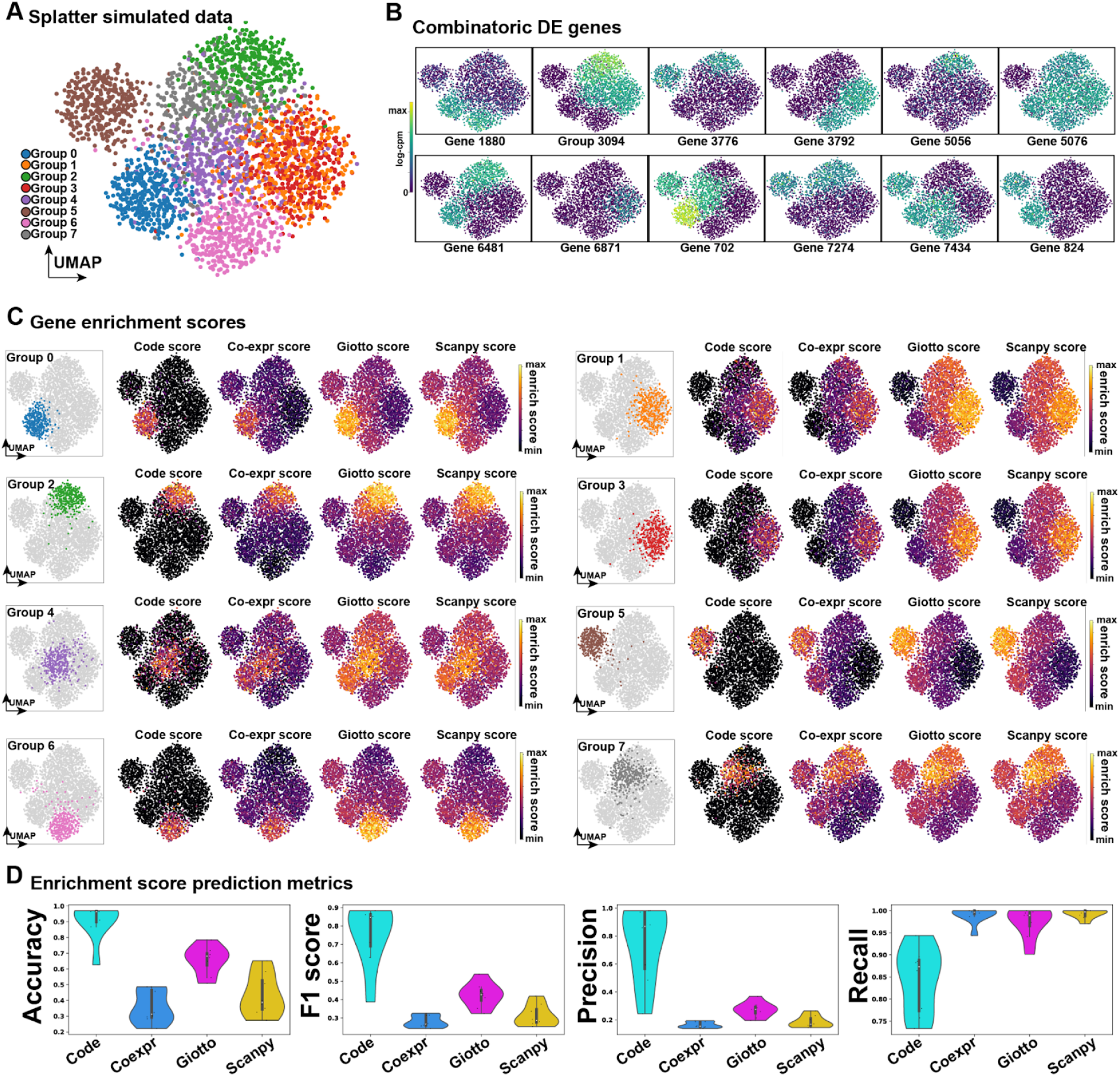
*Cytocipher code-scoring* outperforms existing methods for enrichment of cell population marker gene combinations in simulated data. **A**. Uniform Manifold Approximation (UMAP) of splatter simulated single cell RNA-seq data with 8 groups, each of which has a unique *combination* of gene expression, with single marker genes demarcating a cluster from all other cells only present in a few cases. **B**. UMAPs display the log-counts-per-million for each simulated differential gene between groups, illustrating gene combinatorics for cluster definition. **C**. Results from performing per-cell-gene enrichment using the marker genes for each group illustrated in B. Each sub-panel focuses on a particular group of cells, with the enrichment scores for *Cytocipher code-scoring, Cytocipher coexpr-scoring, Giotto PAGE* enrichment, and *Scanpy-scoring* shown alongside the highlighted cluster; clearly indicating high specificity with minimal background for the *code-scoring* approach. **D**. Enrichment score prediction metrics for predicting each group based on the enrichment scores for each method. Positive scores for each enrichment method were used to indicate cluster membership. Accuracy, F1 score, precision, and recall measures are shown as violin plots, with each point in the violin representing the measure for a given group of cells, and separate violins indicating the enrichment method used for scoring.

We sought to quantitatively test the predictive capability of the scoring methods for binary classification, whereby cells with a positive score for a given method are predicted to belong to the given cluster, and all other cells are considered outside of the cluster. Repeating this for each cluster, we calculated the accuracy, F1 score, precision, and recall of each enrichment method (Figure 2D). Based on these metrics, methods from greatest to lowest performance were *code-scoring, Giotto PAGE, Scanpy* -scoring, and then *coexpr-scoring*. The separation between methods was considerable; *code-scoring* clearly outperformed the alternatives based on all metrics, except for recall, where *code-scoring* was the worst performing. This suggests that the negative gene set filtering can increase false negatives at the single cell level. However, when considering that the F1 score represents the harmonic mean of both the precision and recall^10^, the higher precision of *code-scoring* outweighs the lower recall for better overall binary classification performance in this test case.

Overall, *Cytocipher code-scoring* performed well in the simulated test case, prompting further benchmarking on real data where transcriptionally distinct cells are defined by complex combinatorial gene co-expression.

### *Cytocipher code-scoring* performs comparably to existing methods for binary classification task of E18.5 hypothalamus neuronal subtypes

To assess the utility of *Cytocipher code-scoring* on real data with complex combinatorial gene co-expression, the enrichment scoring methods were applied to our previously published mouse E18.5 hypothalamus scRNA-seq data^11.^ The hypothalamus represents an important test case, as it contains 79 different neuronal subtypes delineated by a small number of marker genes expressed in various unique combinations (Figure 3A). Cluster annotations of the top marker genes from the original publication were used here for gene enrichment analysis (Figure 3A-B). Visualising the per-cell enrichment scores for each cluster (Figure 3B), *Cytocipher code-scoring* and *coexpr-scoring* qualitatively appeared the most specific to cells of each cluster, with minimal cross-cluster scoring evident. However, some clusters for the *code-scoring* appeared to have few cells with positive scores for their respective cluster, while the *coexpr-scoring* did have positive scores. This indicates cases where a given cluster co-expressed genes belonging to the negative set but is not annotated as such. In our previous paper^11^, we manually annotated this data based on the top marker genes following the standard *Scanpy* pipeline^1^, hence highlighting the importance of the *code-scoring* approach for detecting co-expression in complex cases where manual annotation is prone to error without utilisation of a method which can visualise unique combinatorial gene co-expression.

**Figure 3.**
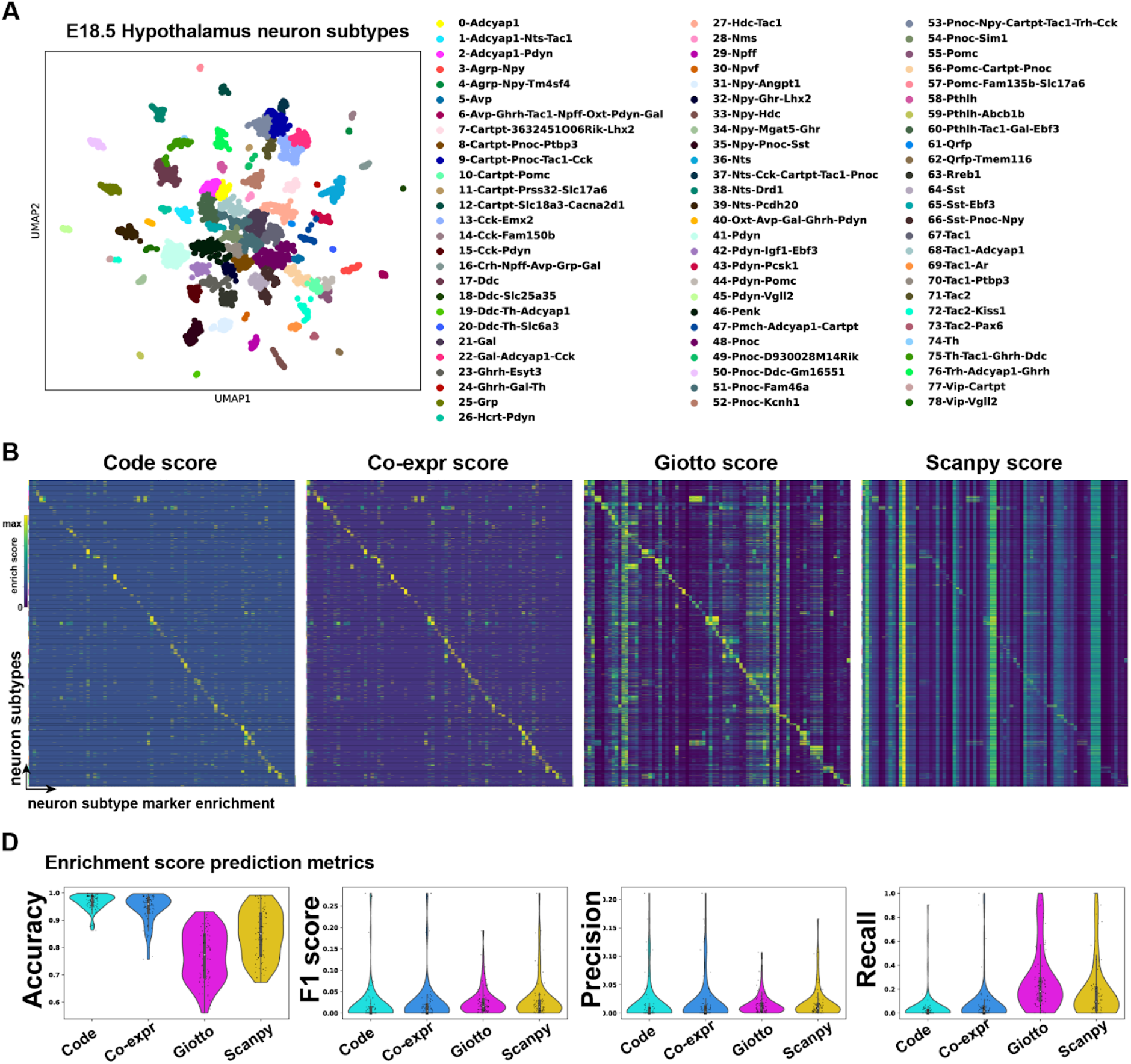
*Cytocipher code-scoring* performs comparably to existing methods for enrichment of cell population marker gene combinations in hypothalamus neuronal subtypes. **A**. UMAP E18.5 hypothalamus single cell RNA-seq depicting 79 neuronal subtypes clusters. **B**. Heatmap displaying per-cell enrichment scores for cluster membership depicted in A; each row is a cell, and each column is a neuronal subtype cluster. Cells and clusters are ordered such that perfect correspondence of cells to score for their respective cluster lies on the diagonal of the heatmap. Hence scoring outside of the diagonal indicates ‘cross-scoring’, where cells also score for gene expression outside of their cluster membership. Lack of scoring along the diagonal indicates cell gene expression does not match cluster membership. **C**. Enrichment score prediction metrics for predicting each neuronal subtype cluster based on the enrichment scores for each method. Positive scores for each enrichment method were used to indicate cluster membership. Accuracy, F1 score, precision, and recall measures are shown as violin plots, with each point in the violin representing the measure for a given neuronal subtype, and separate violins indicating the enrichment method used for scoring.

*Giotto PAGE* enrichment showed considerable cross-clustering scoring, although there is clearly higher enrichment for cells belonging to a cluster than outside of the cluster (Figure 3B). *Scanpy* -scoring on the other hand performed poorly, with considerable cross-cluster scoring and few apparent cases of specificity for scoring cells for their respective clusters.

To quantitatively benchmark the enrichment scoring methods, we again tested using the binary classification task, as in the simulated data example above (Figure 3C). While the *Cytocipher code-scoring* and *coexpr-scoring* give a higher accuracy, F1 was comparable to other methods, with slight improvements in precision. Recall was clearly better for *Giotto PAGE* and *Scanpy* than the *Cytocipher* methods. Overall, while we observed comparable performance between the methods with the F1, there was clearly a lower recall rate but higher precision for the *Cytocipher* methods.

Importantly, *Cytocipher code-scoring* can also be used as a diagnostic tool with which to visualise imperfections in the manual annotations of gene co-expression, and the clear cases of few scores for a given cell type indicate labelling of unique gene co-expression where another cluster must also co-express this combination. This is evidenced by contrasting *code-scoring* with *coexpr-scoring*, where the additional 0 scores in the former indicate cells that expressed genes in the negative gene sets, indicating non-distinct gene expression with atleast one other cluster. Hence in this case, the reduced recall observed is a useful diagnostic with which to assess the uniqueness of cluster marker gene set co-expression at the individual cell level. Prompted by this, we next sought to utilise our enrichment scoring approach to test for significantly different populations of cells in scRNA-seq data.

### *Cytocipher cluster-merge* reveals novel heterogeneity in 3K human peripheral blood mononuclear cells (PBMCs)

To test the capability of *code-scoring* to identify over-clustering in scRNA-seq data and subsequently correct this using *cluster-merge*, we utilised the 3K human PBMC data, a common test dataset for single cell methods due to its well characterised cell types. As shown in the *Scanpy* tutorial^1^, Leiden clustering produced 8 clusters of cells, which could be labelled without any over-clustering or manual merging based upon marker genes to identify B-cells, CD14+ monocytes, Dendritic cells, FCGR3A monocytes, megakaryocytes, CD4+ T cells, CD8+ T cells, and Natural Killer (NK) cells (Figure 4A).

**Figure 4.**
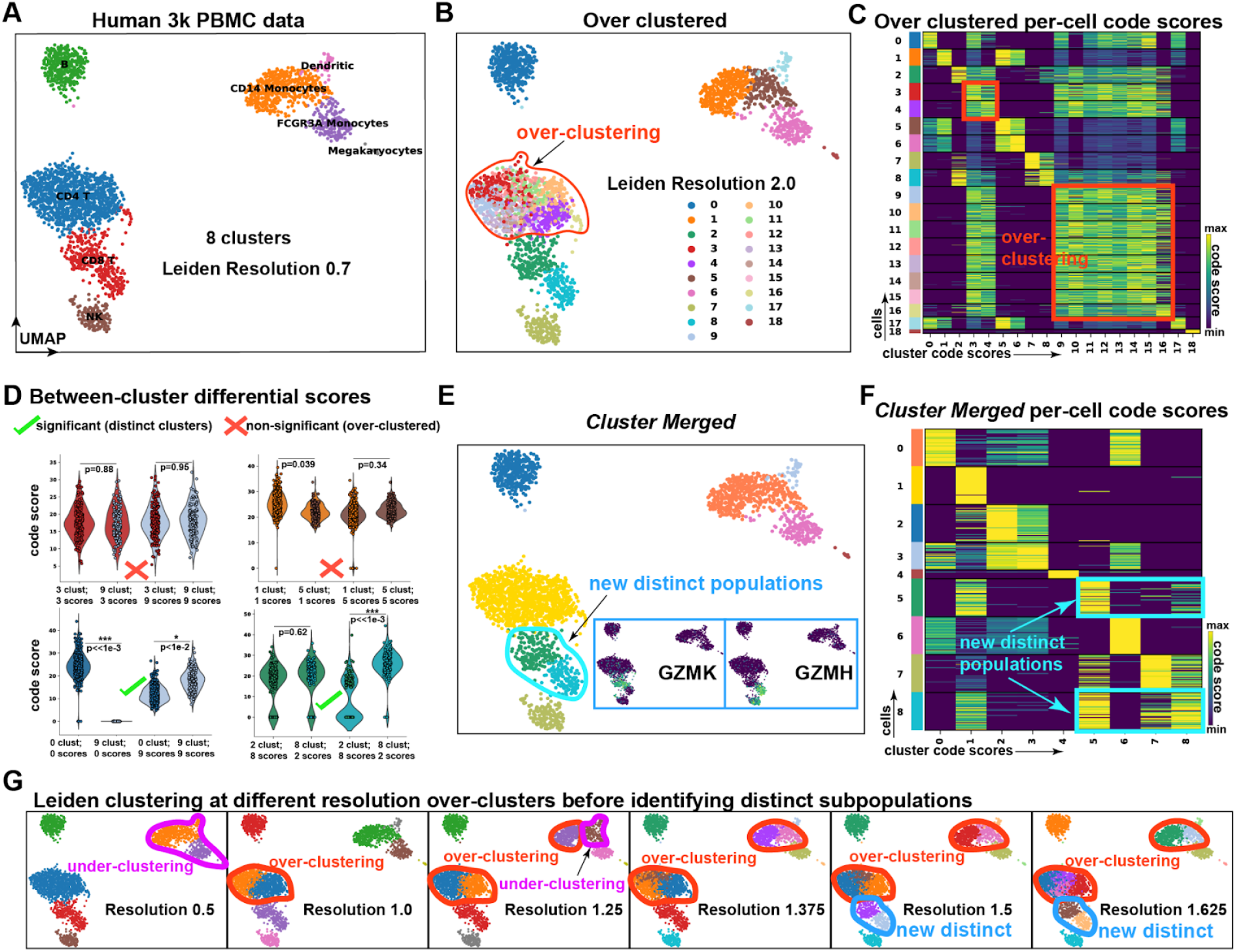
Cross-cluster comparisons of code-scores can be used to merge over-clustered single cell data, revealing novel heterogeneity in human peripheral blood mononuclear cells (PBMCs). **A**. UMAP of Human 3K PBMC scRNA-seq, clustered at Leiden resolution 0.7 to create 8 clusters. Cells are annotated by cell type; B cells, CD4+ T cells, CD8+ T cells, Natural Killer (NK) cells, Dendritic cells, CD14+ Monocytes, FCGR3A+ Monocytes, and Megakaryocytes. **B**. Over-clustering of the PBMC data at Leiden resolution 2.0, producing 19 clusters. **C**. Heatmap depicting *Cytocipher code-scores*, where each row is a cell, and each column is a cluster. Cells and clusters are ordered such that scores along the diagonal indicate scores of cells for their respective cluster. Cross-scoring of cells for different clusters are indicated with red boxes; corresponding to the over-clustering introduced in B. **D**. Violin plots show example cluster significance tests. Non-significant and significant cluster pairs are indicated with red crosses and green ticks, respectively. The y-axes are the code score, and the four violins within each plot indicate combinations of cells belonging to each cluster scoring for their own cluster and the cluster being compared against. Clusters are significantly different when a significant p-value is observed for either set of cluster scores when comparing cells between clusters. **E**. UMAP depicts the PBMCs after merging non-significantly different clusters. New distinct populations are outlined in blue. Small blue boxes of UMAPs depict the log-cpm expression of genes in the new distinct subpopulations (GZMK & GZMH). **F**. The same as C, except for the new clusters after merging. **G**. UMAPs of the data clustered at increasing Leiden resolutions from left to right. Cases of over- & under-clustering are outlined in red and magenta, respectively. The new distinct clusters appear at resolution 1.5 (outlined in blue), at the same point where over-clustering is still clearly evident.

We purposely over-clustered this data, setting a Leiden resolution of 2 to identify 18 clusters, which primarily over-clustered the CD4+ T cells (Figure 4B). *Cytocipher code-scoring*, using automatically identified marker genes clearly indicated this over-clustering, with all of the CD4+ T cell subclusters clearly all cross-scoring with one another (Figure 4C). Importantly, using the previous PBMC cell type annotations as the ground-truth (Figure 4A) to determine non-significantly different clusters in the over-clustered cells (Figure 4A-B), we tested a range of possible implementations for particular design choices for cluster pair significance testing within *Cytocipher*, measuring test performance and bias in test significance (Figure S1; see Methods). The best implementation significantly de-correlated test significance with cell pair abundance (Spearman’s *ρ*=0.083) and had high test performance (AUROC=0.9803), thus prompting this implementation as the default setting for *Cytocipher cluster-merge* (see Methods for details).

We subsequently applied *Cytocipher cluster-merge* with the default settings on this overclustered data, visualising the distributions of the code-scores using the combination of cluster profiles and cells in each cluster to show significant versus non-significant clusters (Figure 4D). This approach correctly identified no significant difference between clusters 3 and 9, nor clusters 1 and 5 (Figure 4D); the former are over-clusters of the CD14+ monocytes, while the latter are over-clusters of the CD4+ T-cells (Figure 4A-B). Clusters 0 and 9, representing B cells and dendritic cells, were correctly identified as different (Figure 4D, A-B). Interestingly, all clusters were correctly merged after one iteration to the ground-truth cell types, except clusters 2 and 8; subclusters of the CD8+ T cells (Figure 4D). Note that after the merge operation cluster 2 was relabelled to cluster 5 (Figure 4E). Inspection of the marker genes for these clusters revealed specific expression of GZMK in merged cluster 5, and GZMH in merged cluster 8 (Figure 4E). GZMK+ CD8+ T cells were recently identified as an important CD8+ T cell subtype in ageing mice, where specific expansion of these cell types is associated with inflammation, thought to be caused by increased secretion of the protein of GZMK^17^. GZMH, on the other hand, has been used as a marker to identify activated cytotoxic effector CD8+ T cells in contrast with transitional CD8+ T cells^18^.

*Cytocipher cluster-merge* was able to detect additional heterogeneity in CD8+ T cells, not previously described in this dataset (Figure 4E-F). We tested whether this division of CD8+ T cells was detectable at a Leiden resolution without also over-clustering the data (Figure 4G). At Leiden resolution 0.5, we identified under-clustering of the CD14+ monocytes and dendritic cells. Resolution 1 corrected this under-clustering but resulted in over-clustering of the CD4+ T cells. Resolution 1.25 mis-clustered the dendritic and CD14+ monocytes, with the over-clustering of the CD4+ T cells retained. We first observed the new distinct subdivision of CD8+ T cells at resolution 1.5, at the same time there is over-clustering of the CD4+ T cells and CD14+ monocytes (Figure 4G).

While it was not possible to exhaustively test every possible resolution, we could not find a resolution where the CD8+ T cell subpopulations were identifiable using Leiden without also over-clustering. Hence, while *Cytocipher cluster-merge* automatically identified these important subpopulations, Leiden analysis could not detect these subpopulations without over-clustering other cell types.

### *Cytocipher cluster-merge* identifies intermediate states in pancreas development

In the case of scRNA-seq analysis of developmental data, intermediate cell states can exist as a continuum of cells between progenitor cells and mature cell types^19^. The continuity of developmental scRNA-seq makes determination of discrete populations with standard clustering ambiguous, since clear separation of cell populations is not evident. Given that *Cytocipher* considers gene expression combinations for scoring cluster membership, we hypothesised that *Cytocipher* would be sensitive to transcriptionally distinct intermediate states, potentially allowing for identification of fine-grained branch points that represent lineage decisions toward terminal cell fates.

To test this hypothesis, we utilised scRNA-seq data from mouse E15.5 pancreas development^16^, since this has been extensively studied with the tool *Scvelo*^*12*^, an analysis method for predicting the direction of cell differentiation. We first reproduced *Scvelo* predictions of cell differentiation in the pancreas data^12^ (Figure 5A). Over-clustering the pancreas data to 30 clusters, and subsequently applying *Cytocipher code-scoring* showed considerable cross-scoring between clusters (Figure 5B). Importantly, the *code-scoring* approach was still highly binary in the case of continuous developmental clusters; as evidenced by the lack of low-level background cross-scoring, allowing for clear identification of over-clusters (Figure 5B).

**Figure 5.**
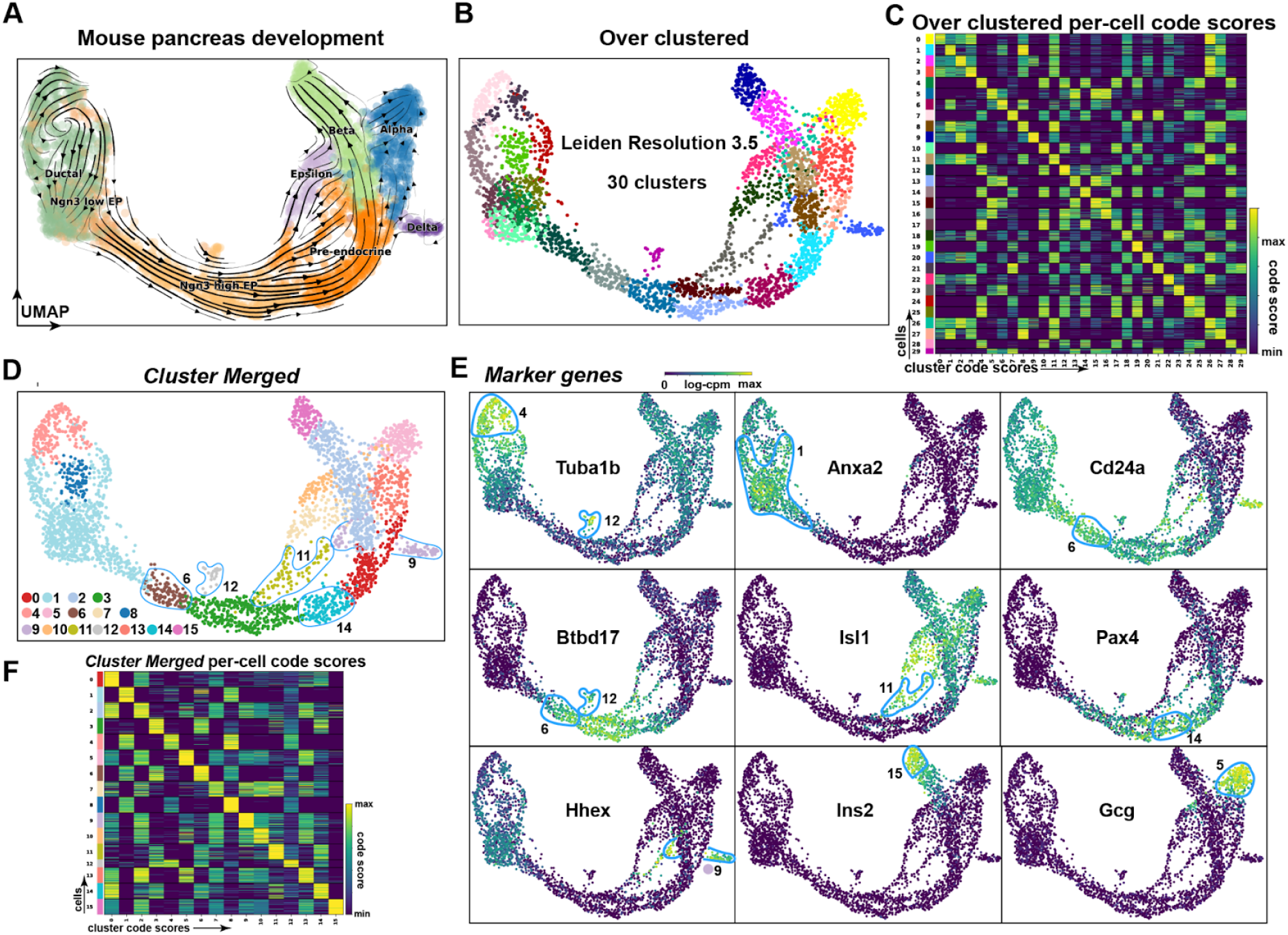
*Cytocipher* identifies intermediate cell states in mouse pancreas development. **A**. UMAP of mouse E15.5 pancreas cells, with cells annotated by broad cell types; ductal, Ngn3 low endocrine progenitor cells (EP), Ngn3 high EP cells, pre-endocrine, epsilon, beta, alpha, and delta. **B**. UMAP with cells over-clustered at Leiden resolution 3.5, producing 30 clusters. **C**. Heatmap of *Cytocipher* code-scores per cluster, as shown for clusters in panel **B. D**. UMAP of 16 significant clusters determined from *Cytocipher cluster-merge* applied to the clusters depicted in panel B. Novel intermediate states outlined. **E**. Top marker genes determined for the clusters depicted in D, with cells belonging to the relevant clusters of the marker genes outlined (see text for interpretation). **F**. Heatmap of *Cytocipher* code-scores per cell (row) and cluster (column), for merged clusters depicted in panel D.

Applying *cluster-merge* resulted in 16 distinctly different clusters, including several intermediate cell states not identified with the original annotations (Figure 5A,C, F). We then examined the marker genes automatically determined by *Cytocipher* after cluster merging, which revealed that the intermediate states identified expressed several important regulators of pancreas development (Figure 5E). For instance, cluster 6 clearly grouped an intermediate state transition from early endocrine progenitor (EP) cells, primarily expressing Anxa2, to Btbd17 expressing EP cells; a known marker of Ephi EP cells^20^ (Figure 5D-E). Clusters 11 and 14 clearly highlighted alternate lineage paths, with the former expressing Isl1 and the latter Pax4, both important developmental transcription factors controlling pancreas development^21,22^ (Figure 5D). Importantly, Pax4 has been previously shown as a lineage determinant for pancreatic beta cell fate^22^, and, inline with *Scvelo* differentiation predictions, cluster 14 appeared to represent an early decision toward this cell type not identified in the original annotations (Figure 5A). Importantly, *Cytocipher* cluster 9 appeared to branch off cluster 11 (Figure 5D), with cluster 9 cells expressing Hhex, a homeodomain transcription factor required for pancreatic delta cell determination^23^ (Figure 5E).

*Cytocipher* was able to identify potentially important intermediate states in developmental scRNA-seq data, evident when examining the outputted marker genes for the determined intermediate lineage states, which included several well-known transcription factors required for determination of the cell fates highlighted (Figure 5D-E).

### *Cytocipher* detects over-represented subpopulations in prostate cancer

Comparison between normal and tumour scRNA-seq samples can reveal over- and under-represented cell types in cancer, indicating important cell types that may contribute to cancer progression and therapeutic resistance. We hypothesised that application of *Cytocipher* within a cancer context would identify cancer relevant cell subpopulations and the key genes defining these populations. To test this hypothesis, we compared *Cytocipher* significant clusters in the context of prostate cancer to significantly differentially abundant (DA) cell populations independently determined with *Milo*^*14*^ (Figure 6); a novel DA analysis method that considers sample information and the cell-cell neighbourhood graph to determine small populations of cells differentially abundant between two or more conditions^14^.

**Figure 6.**
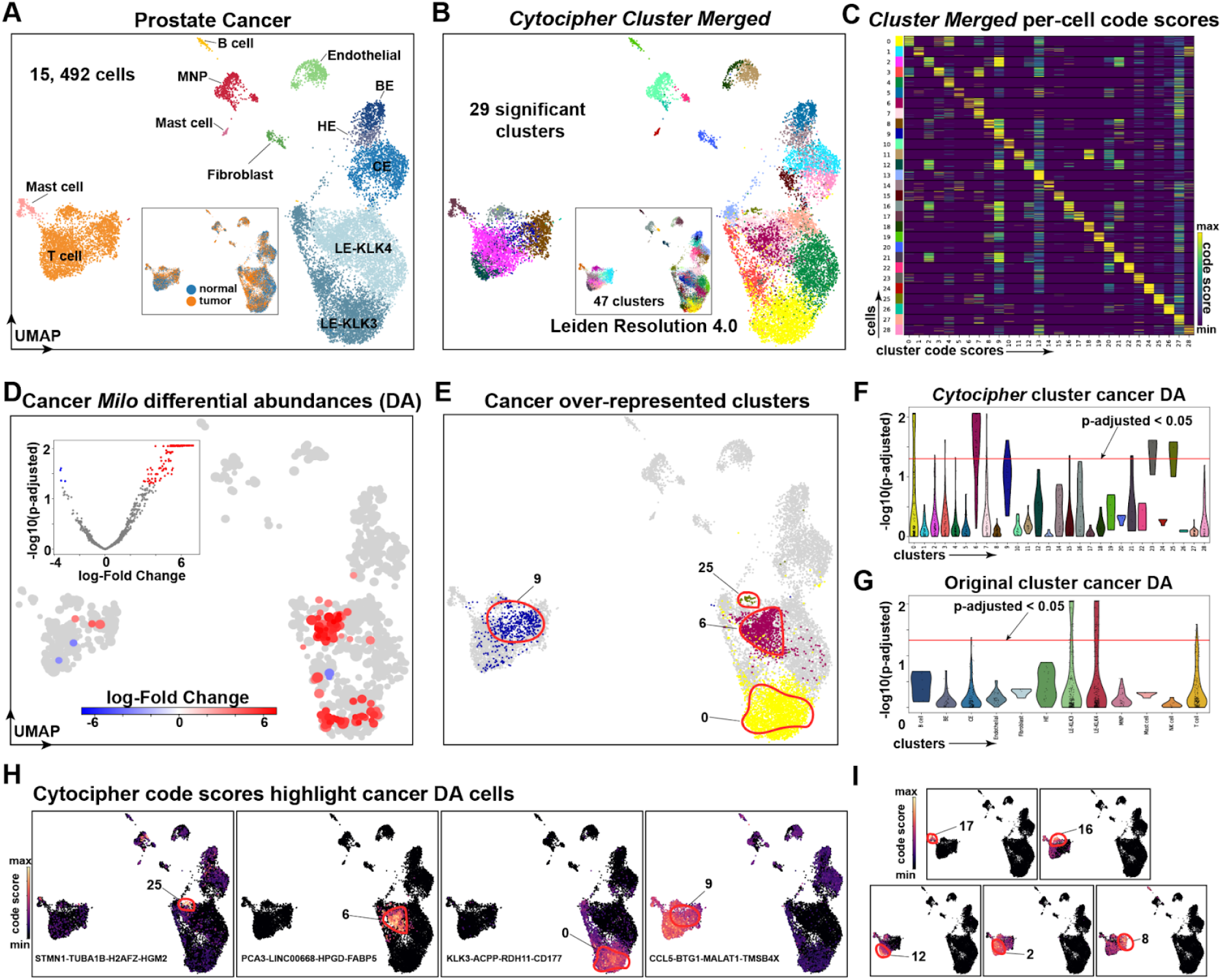
*Cytocipher* detects over-represented sub-populations in prostate cancer. **A**. UMAP of scRNA-seq from normal and tumor tissue from the human prostate, consisting of 15,492 cells. Data as provided by Tuong, *et al*. (2021). **B**. *Cytocipher* significant clusters after merging 47 clusters produced from Leiden clustering at resolution 4.0 (depicted as inner UMAP). **C**. Heatmap depicting *Cytocipher* code-scores, where each row is a cell, and each column is a cluster. Cells and clusters are ordered such that scores along the diagonal indicate scores of cells for their respective cluster. Code-scores for the 29 significant clusters in panel B are shown. **D**. Inner volcano plot depicts -log10(p-adjusted) on the y-axis and log-fold change on the x-axis testing for differential abundance of cells on the cell-cell neighbourhood graph using *Milo*. The outer UMAP depicts non-significant cells in grey, and significantly differentially abundant (DA) cellular neighbourhoods in red (over-represented in prostate cancer) and blue (under-represented in prostate cancer). **E**. UMAP highlights the significant clusters detected by *Cytocipher* that were independently determined as over-represented in prostate cancer by *Milo* DA analysis. **F**. Violin plots with -log10(p-adjusted) on the y-axis and *Cytocipher* significant clusters on the x-axis. A red horizontal line indicates the p-adjusted cutoff of 0.05, cells above the red-line are considered significantly DA between tumor and cancer samples. **G**. Equivalent to F, except for the original prostate cell types depicted in A. **H**. *Cytocipher* code-scores for the prostate cancer over-represented clusters detected by *Cytocipher*, from left-to-right code-scores depicted are specific to cluster 25, 6, 0, and 9. The marker gene set for each respective cluster is shown within the UMAP plots. **I**. Equivalent to H, except depicting clusters 17, 16, 12, 2, and 8. These clusters also score for cluster 9 as depicted in H, but are significantly different due to additional gene co-expression.

The prostate cancer scRNA-seq dataset included integrated healthy and cancerous prostate tissue representing 15,492 cells from 10 patients and 24 samples^13^ (Figure 6A). 12 cell types are annotated by the original authors, with prostate specific cell types including basal (BE), hillock (HE), club cells (CE), and two luminal epithelial cell types (LE-KLK3 and LE-KLK4)^13^ (Figure 6A).

We over-clustered the prostate cancer scRNA-seq data using Leiden clustering at resolution 4.0 to produce 47 clusters, from which 29 significantly different clusters were determined using *Cytocipher cluster-merge* (Figure 6B). Examination of the *Cytocipher* code-scores per significant cluster indicated highly specific marker gene co-expression differentiated each cluster (Figure 6C). Independent application of *Milo* revealed 4 populations of over-represented cells in the prostate cancer samples, all of which tightly corresponded to significant clusters identified by *Cytocipher* but not by the original annotations (Figure 6D-G). In line with the original publication and the prostate cancer literature^13,24^, three DA populations were within the luminal epithelial cells (Figure 6A,E). The remaining population represented a T cell subpopulation (Figure 6A,E).

Highly specific *Cytocipher* code-scores highlighted each of the DA luminal epithelial cells (Figure 6H). *Cytocipher* marker genes used for *code-scoring* for each of the DA luminal epithelial cells were also highly relevant for prostate cancer. For example, cluster 25 included STMN1 as a marker gene, a known over-expressed oncoprotein in prostate cancer that controls cell proliferation^25^ (Figure 6H). Cluster 6 included PCA3, HPGD, and FABP5; the former produces a protein which is a key prostate cancer biomarker in urine tests^26^, while the latter two genes have been implicated as metabolic regulators of prostate cancer^27,28^. The third significant luminal epithelial subpopulation (cluster 0) had marker genes which included ACPP, the gene encoding the prostate cancer biomarker PAP^29^.

The *Milo* DA T cell subpopulation, which was also detected by *Cytocipher* (cluster 9), did not express highly specific code-scores (Figure 6H). Cluster 9 was however distinct from other *Cytocipher* clusters by *not* scoring for marker genes of other T cell subsets (Figure 6I). This emphasises the importance of the bi-directional test utilised by *Cytocipher cluster-merge*, which requires subpopulations to be *mutually* non-significantly different for the cluster pair to be merged (Figure 1B3).

Overall, these findings demonstrated that *Cytocipher* can produce cancer relevant sub-populations defined by biologically relevant genes, such as cancer biomarkers and regulators.

### *Cytocipher* analysis of >480,000 cells in Tabula Sapiens Atlas reveals scalability and high-test performance

Atlas scale scRNA-seq data can contain hundreds of cell types and exceed hundreds of thousands of cells. Analysing such data requires highly scalable and accurate methods that are also time and memory efficient. We tested the scalability and performance of *Cytocipher* by analysing the Tabula Sapiens Atlas^3^, which consists of 483,152 cells across 24 tissues and 177 cell types^3^ (Figure 7A). The cell type annotations were manually annotated by a team of experts for each tissue type^3^, thereby presenting an extremely high quality ground-truth of cell types to test *Cytocipher*. First, we applied *Cytocipher code-scoring* on the 177 cell types to confirm annotation quality (Figure 7B). This revealed high correspondence between cell type annotations and cell type code-scores; confirming the 177 manual annotations were transcriptionally distinct (Figure 7B). Treating the high-quality Tabula Sapiens annotations as a ground-truth, we then randomly sampled within each cell type to artificially split cell types into 3 artificial clusters (see Methods). This gave ground-truth over-clusters consisting of 503 cell populations with which to test method performance, whereby a pair of artificial clusters occurring *within* a cell type were considered *non-significantly* different and over-clusters from *different* cell types were considered *significantly* different. Application of *Cytocipher cluster-merge* to these artificial over-clusters revealed some over-merging, with the top 4 most over-merged clusters consisting of artificial clusters both within- and between-cell type annotations (Figure 7C). However, closer examination of these over-merged cases revealed shared biology, such that fine-grained cell types had been merged. For example, smooth muscle cells from different parts of the body had been merged, such as vascular and bronchial smooth muscle (Figure 7C). Different T cell subtypes were also merged to a single cluster. CD4+ and CD8+ T cells were detected however, and their subtypes merged into independent CD4+ and CD8+ clusters (Figure 7C). Thus, while *Cytocipher* did not perform perfectly per the ground truth, related function amongst the cell types merged was observed.

**Figure 7.**
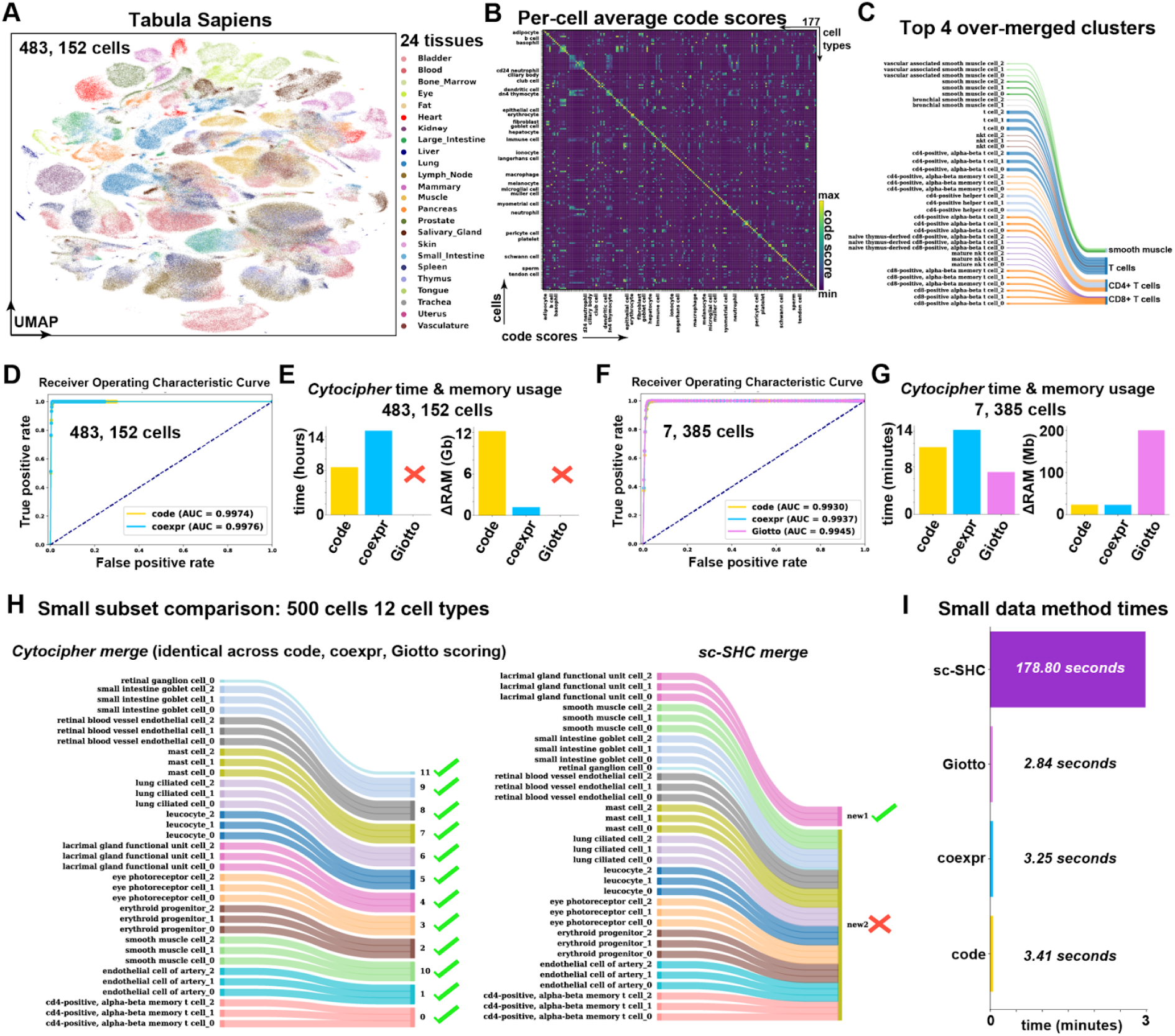
*Cytocipher* scales to >480,000 cells with high test performance. **A**. Tabula Sapiens UMAP depicting 483,152 cells sampled from 24 tissues by *The Tabula Sapien Consortium* (2022) **B**. Heatmap depicting *Cytocipher* code-scores for the 177 cell types annotated within the Tabula Sapiens dataset. Each row is a cell, and each column is a cluster. Cells and clusters are ordered such that scores along the diagonal indicate scores of cells for their respective cluster. **C**. Sankey diagram, indicating the top 4 over-merged clusters by *Cytocipher* when testing with artificial random sub-groups of the 177 cell types. The left side of the diagram indicates the sub-grouped cell types, and the right side indicates the sub-grouped cell types merged by *Cytocipher*. **D**. Receiver operating characteristic (ROC) curve depicting the true-positive-rate on the y-axis and the false-positive-rate on the x-axis at different p-value cutoffs using *Cytocipher cluster-merge* applied to artificial random subgroups of the 177 cell types using either *code-scoring* or *coexpr-scoring*. Area under the curve (AUC) for each scoring method is indicated in the legend. **E**. Bar charts indicate time and memory usage of Cytocipher when analysing the 483,152 cells across artificial cell type sub-groups. Giotto scoring could not be performed on the full dataset due to memory limitations. **F-G**. Equivalent to **D-E**, except downsampling each of the cell type subgroups to a maximum of 15 cells to reduce the dataset size to 7,385 cells, enabling *Giotto-scoring* to be run for comparison and examine the effect of fewer cells on test performance. **H**. Small comparison between *Cytocipher* and *Sc-SHC*, using the 500 cells and 12 random cell types subsetted from the 177 cell types. Green ticks on the right hand side of the Sankey diagrams indicate artificial over-clusters were correctly merged, while red crosses indicate incorrect merging. **I**. Bar plot indicating run-time for the different methods, with methods depicted on the y-axis and run-time on the x-axis.

We then performed a systematic test of *Cytocipher* performance, examining how different p-value cut-offs affected the true positive and false positive rates for classifying significant versus non-significantly different cluster pairs using the aforementioned ground-truth (Figure 7D). ROC curves were generated from applying *Cytocipher cluster-merge* with both *code-scoring* and *coexpr-scoring*. AUROC was used to quantify test performance, with an AUROC of 1 indicating perfect ability to discriminate significant versus non-significant cluster pairs and 0.5 as random classification (Figure 7D). *Cytocipher cluster-merge* with both *code-scoring* and *coexpr-scoring* achieved almost perfect AUROC values; 0.9974 for the former and 0.9976 for the latter (Figure 7D). Time and memory requirements for this task were also measured for the two different scoring methods. *Cluster-merge* with *code-scoring* took 8.5 hours to compute, while *coexpr-scoring* was 15 hours. Memory requirements were an additional 12.2 giga-bytes (Gb) of RAM for *code-scoring* and 1.19 Gb for *coexpr-scoring*. We were not able to run *Giotto-scoring* because it exceeded our available RAM capacity.

Due to the high memory requirements for *Giotto-scoring*, we then tested the effect of subsetting the data. Downsampling to a maximum of 15 cells for each one of the 503 overclusters resulted in a total of 7,385 cells. Performing the same benchmarking as with the full dataset, and including *Giotto-scoring* due to the lower RAM requirements with the reduced data set, revealed greatly reduced time and memory performance with negligible effects on test performance (Figure 7F-G). *Cytocipher cluster-merge* applied with *code-scoring, coexpr-scoring*, and *Giotto-scoring* all produced high AUROC values; 0.9930, 0.9937, and 0.9945, respectively (Figure 7F). Run time was also reduced from hours to <15 minutes in each case (Figure 7G). Memory usage was reduced from Gb to mega-bytes (Mb), with Giotto having a 10-fold higher memory requirement of 200 Mb, while *code-scoring* and *coexpr-scoring* peaked at 23.7 Mb and 23.39 Mb, respectively. This demonstrates that downsampling of over-clusters is an effective strategy for determining significantly different clusters with *Cytocipher* while reducing compute requirements. Overall, we found that *Cytocipher* scales to large atlas data with >480,000 cells and hundreds of cell types with high performance.

While *Cytocipher* represents the first method for cluster significance analysis in scRNA-seq data, during the preparation of this manuscript another distinctly different method, *Single cell Significance of Hierarchical Clustering* (*Sc-SHC*), was publicly released^31^. *Sc-SHC* differs from *Cytocipher* in the usage of raw count data, Poisson count modelling, Silhouette scoring, and utilisation of a dendrogram to test branch-points for significantly different groups of clusters rather than cluster pairs^31^. Further contrasting with *Cytocipher, Sc-SHC* does not include the output of marker gene sets which significantly differentiate each cluster. We compared run-time and performance between *Cytocipher* and *Sc-SHC* in a small test case, where 12 cell types were randomly sampled from the 177 cell types in the downsampled Tabula Sapiens data, resulting in 500 cells (Figure 7H-I). Application of *Cytocipher cluster-merge* with *code-scoring, coexpr-scoring*, and *Giotto-scoring* produced the same result; correctly merging all the artificial over-clusters to the original cell types (Figure 7H). *Sc-SHC* correctly merged lacrimal gland functional unit cell over-clusters, but incorrectly merged all other cell types (Figure 7H). *Cytocipher cluster-merge* was also significantly faster, taking <3.5 seconds on the 500 cells and 34 over-clusters, while *Sc-SHC* had a 178.80 second compute time (Figure 7I). Overall, this test suggested *Cytocipher* cluster significance analysis is more accurate and time efficient than the concurrently developed method *Sc-SHC*.

## Discussion

ScRNA-seq is having a transformative effect upon our understanding of cellular diversity in multicellular organisms, and several technological advances are resulting in an extensive build-up of data. However, the bioinformatics analysis of scRNA-seq data remains challenging and involves problematic post host manual curation. *Cytocipher* represents the first method that performs statistical analysis of tentative single cell clusters to ensure that cell groups represent significantly transcriptionally distinct populations. In contrast to standard scRNA-seq clustering, which operate on the nearest-neighbour graph, *Cytocipher* refers back to the original gene expression measurements, and performs per-cell enrichment scoring for cluster marker genes and a bi-directional statistical test to infer significantly different clusters. In this study, we demonstrate that this statistical approach ensures the final single cell clusters align with transcriptionally distinct populations of cells allowing for improved insights in various biological contexts.

When analysing the human PBMC data, we did not find any Leiden resolution that could accurately uncover all of the transcriptionally distinct cell populations without also over-clustering. However, *Cytocipher* was able to identify the transcriptionally distinct populations. Importantly, several methods have been developed concerned with varying cluster hyper-parameters to identify an optimal parameter set^32–39^. While the criteria to define ‘optimal’ clustering is tool dependent^2–39^, the tool *Clustree*^*39*^ for example, defines this as the highest resolution clustering where further increasing the clustering resolution results in arbitrary rearrangements of cells to increase the number of clusters. All of these approaches pre-supposes that there exists a set of hyper-parameters where the clustering method utilised can identify the underlying transcriptionally distinct cell populations, despite this not being a part of the objective function utilised by current clustering methods. Our results suggest that future methods should instead evaluate single cell clustering by determining if tentative clusters are supported by cells within the clusters displaying distinctly different gene co-expression. This approach improves interpretability and is more in-line with the goal of scRNA-seq analysis for cell type discovery.

*Cytocipher* considers cluster significance analysis at the level of cluster pairs. An alternative approach, as utilised by the concurrently developed *Sc-SHC*^*31*^, is to use a dendrogram to group clusters and test the branch points to compare groups of clusters. When comparing *Cytocipher* and *Sc-SHC*, we observed the former correctly grouped cell types while the latter incorrectly merged several distinctly different cell types. The improved performance of *Cytocipher* could be due to an averaging effect within *Sc-SHC* when comparing cluster groups. For instance, when *Sc-SHC* tests a particular branch point representing groups of clusters, on average most of the clusters may not be significantly different from one another resulting in a call of non-significant difference for all grouped clusters. However, when *Cytocipher* performs comparisons at the level of cluster pairs, subsets of clusters are identified as distinctly different. To confirm this, systematic testing of every methodological difference between the two methods would be required, which is outside of the scope of the current study. Based on our current results, we suggest the choice of cluster pairs as the unit of testing for future methods of cluster significance analysis to avoid potential averaging effects.

One limitation of *Cytocipher* is its dependency on the cell-cell neighbourhood graph indirectly by the input of Leiden clusters. Because the graph is constructed based upon linearly reduced dimensions from highly variable genes, the clusters tested by *Cytocipher* will define transcriptionally distinct cell populations that are biased towards the highly variable genes used to construct the original neighbourhood graph. It therefore remains an interesting and open question whether alteration of the gene sets utilised to define the initial clusters has an effect on which cells are significantly different from one another; particular in the case of neurons, where several different biological features are used to define neuronal subtypes in different contexts^40^. In summary, *Cytocipher* represents an important advance in the analysis of scRNA-seq data and will enable the reproducible and statistically significant identification of single cell populations that have distinct gene co-expression. In the future, we anticipate similar implementations of cluster significance analysis could be utilised to analyse other complex and heterogeneous biological data such as spatial RNA-seq and scATAC-seq.

## Acknowledgements

We thank Gabriel Foley, Samuel Davis, Sanjana Tule, Ariane Mora, and Samuel Granjeaud for critically reading this manuscript and providing valuable discussion/feedback.

---

**Sup. Figure 1.**
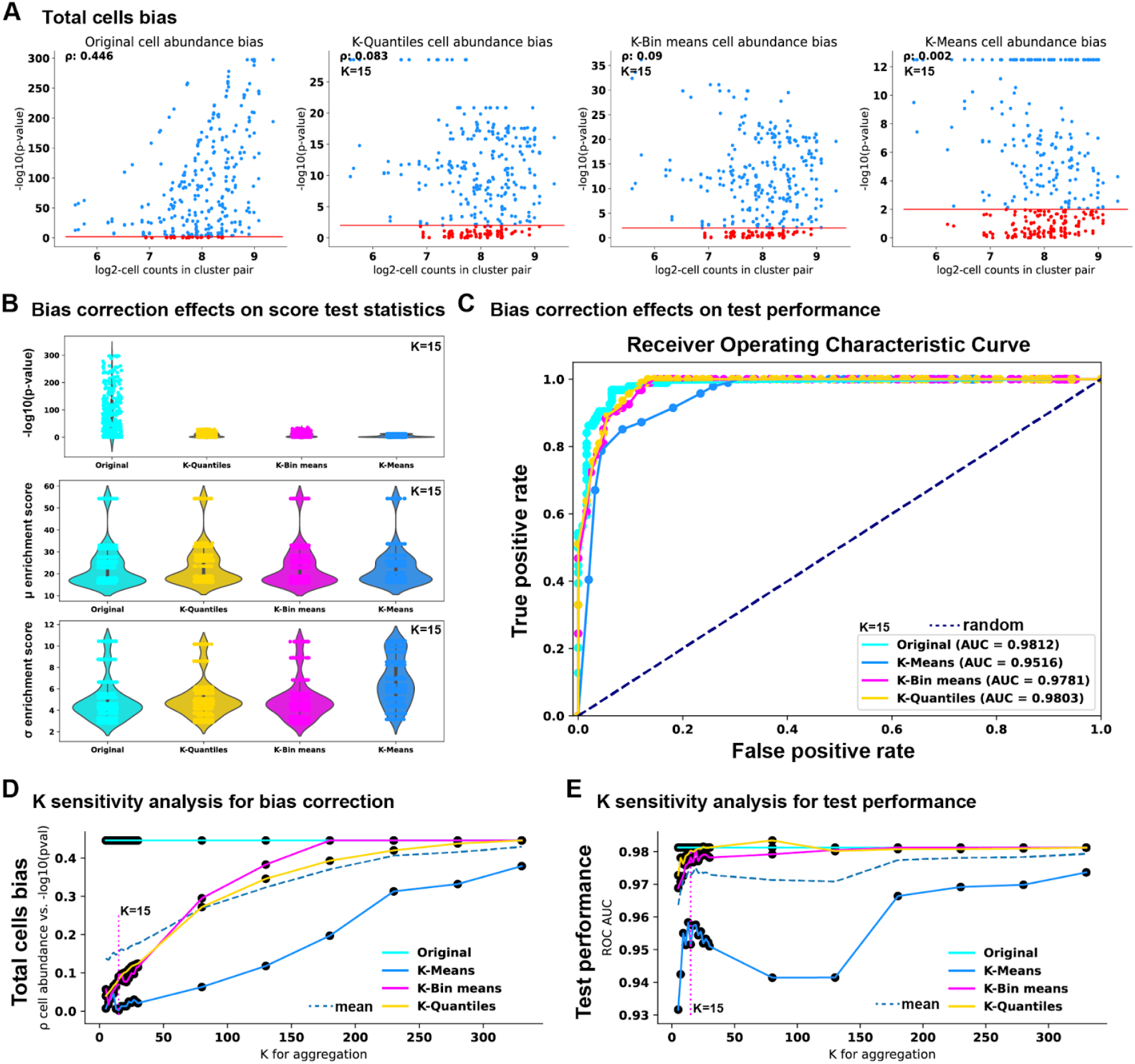
*Cytocipher* code-score summarisation prior to testing corrects significance bias toward higher cell abundance while maintaining high test performance. **A**. Scatter plots depicting -log10(p-value) and log2 of total cell counts in each cluster pair using over-clustered (Leiden resolution 2.0) peripheral blood mononuclear cells (PBMCs). Each individual point represents a pair of clusters which were compared for significant differences using code-scores. Red points are significantly different cluster pairs at p=0.01, with the red vertical line indicating this cutoff. ρ depicts Spearman’s correlation for cluster pair significance and the total number of cells in each cluster (the y- and x-axes, respectively). Separate scatter plots represent the same test applied with different methods for summarising enrichment scores prior to significance testing. From left to right, no summarisation (original), K-Quantiles summarisation, K-Bins summarisation, and summarisation by K-Means are shown. K=15 was used in each case. Each method successfully reduced the correlation of cluster pair significance (-log10(p-values)) and the total number of cells. **B**. Violin plots depict the effect of summarisation methods on key test statistics; -log10(p-values) and the mean/standard deviations of code enrichment scores within each cluster. All methods reduced the p-value inflation while maintaining the mean and standard deviation of the enrichment scores, with the exception of K-Means which inflated the standard deviations. **C**. Receiver operating characteristic (ROC) curve benchmarking each method. Legend indicates area under the curve (AUC) scores (1 indicates perfect classification). K-Quantiles had the best test performance (AUC=0.9803), prompting this as the default summarisation method prior to *Cytocipher* cluster pair significance testing. **D**. Line plot displaying sensitivity analysis when varying the K-parameter and measuring the effect on cell abundance bias. Magenta dotted line indicates K=15 value. Cell abundance bias increased with *k*. **E**. Line plot for K-parameter sensitivity with respect to ROC AUC scores. Test performance was marginally affected in the range *k*>15. *k*<15 resulted in a drop in performance. This prompted k=15 as the default.

